# Controlling the Growth of the Skin Commensal *Staphylococcus epidermidis* Using D-Alanine Auxotrophy

**DOI:** 10.1101/2020.03.10.985911

**Authors:** David Dodds, Jeffrey L. Bose, Ming-De Deng, Gilles Dubé, Trudy Grossman, Alaina Kaiser, Kashmira Kulkarni, Roger Leger, Sara Mootien-Boyd, Azim Munivar, Julia Oh, Matthew Pestrak, Komal Rajpura, Alex Tikhonov, Traci Turecek, Travis Whitfill

## Abstract

Using live microbes as therapeutic candidates is a strategy that has gained traction across multiple therapeutic areas. In the skin, commensal microorganisms play a crucial role in maintaining skin barrier function, homeostasis, and cutaneous immunity. Alterations of the homeostatic skin microbiome are associated with a number of skin diseases. Here, we present the design of an engineered commensal organism, *Staphylococcus epidermidis*, for use as a live biotherapeutic product (LBP) candidate for skin diseases. The development of novel bacterial strains whose growth can be controlled without the use of antibiotics, or genetic elements conferring antibiotic resistance, enables modulation of therapeutic exposure and improves safety. We therefore constructed an auxotrophic strain of *S. epidermidis* that requires exogenously supplied D-alanine. The *S. epidermidis* strain, NRRL B-4268 Δ*alr1*Δ*alr2*Δ*dat* (SE_ΔΔΔ_) contains deletions of three biosynthetic genes: two alanine racemase genes, *alr1* and *alr2* (SE1674 and SE1079), and the D-alanine aminotransferase gene, *dat* (SE1423). These three deletions restricted growth in D-alanine deficient media, pooled human blood, and skin. In the presence of D-alanine, SE_ΔΔΔ_ colonized and increased expression of human β-defensin 2 in cultured human skin models *in vitro*. SE_ΔΔΔ_, showed a low propensity to revert to D-alanine prototrophy, and did not form biofilms on plastic *in vitro*. These studies support the potential safety and utility of SE_ΔΔΔ_ as a live biotherapeutic strain whose growth can be controlled by D-alanine.

## INTRODUCTION

Commensal microorganisms play a crucial role in maintaining human health across a number of organ systems, particularly in the skin (Wikoff, Anfora et al. 2009, Salzman, Hung et al. 2010, Diaz Heijtz, Wang et al. 2011, Grice and Segre 2011, Kau, Ahern et al. 2011, Ravel, Gajer et al. 2011, Mulcahy, Geoghegan et al. 2012). Diverse communities of microorganisms populate the skin, and a square centimeter can contain up to a billion microorganisms (Weyrich, Dixit et al. 2015). These diverse communities of bacteria, fungi, mites and viruses can provide protection against disease and form dynamic, yet distinct niches on the skin (Oh, Byrd et al. 2014). Increasing evidence has associated altered microbial communities or dysbiosis in the skin with cutaneous diseases (Oh, Freeman et al. 2013, Weyrich, Dixit et al. 2015), especially atopic dermatitis (Grice 2014, Powers, McShane et al. 2015, Wan and Chen 2020). New strategies have emerged using microbes as therapy for treating a range of diseases (Dreher-Lesnick, Stibitz et al. 2017). While this approach has gained particular attention in developing biotherapeutics for gastrointestinal diseases (Dore, Multon et al. 2017, Vemuri, Gundamaraju et al. 2017), few studies have reported on using live microbes to treat skin diseases.

*Staphylococcus epidermidis*, recently dubbed as the “microbial guardian of skin health” (Stacy and Belkaid 2019), is a strong candidate for use as a live biotherapeutic product (LBP) for skin conditions*. S. epidermidis* is a Gram-positive bacterium that is ubiquitous in the human skin and mucosal flora. As one of the earliest colonizers of the skin after birth, *S. epidermidis* plays an important role in cutaneous immunity and maintaining microbial community homeostasis (Naik, Bouladoux et al. 2015, Linehan, Harrison et al. 2018). In the clinical setting, *S. epidermidis* has demonstrated activity as a potential therapeutic (Iwase, Uehara et al. 2010, Nodake, Matsumoto et al. 2015, Nakatsuji, Chen et al. 2018). In Japan, a double-blind, randomized clinical trial, topical application of autologous *S. epidermidis* in healthy volunteers increased lipid content of the skin, suppressed water evaporation and improved skin moisture retention while showing no signs of erythema (Nodake, Matsumoto et al. 2015). Others have shown that *S. epidermidis* is capable of producing antimicrobial peptides (AMPs) that selectively target *Staphylococcus aureus* (Cogen, Yamasaki et al. 2010, Cogen, Yamasaki et al. 2010). *S. epidermidis* has also shown other potent, therapeutic effects in preclinical settings. For example, a recent report described anti-neoplastic properties of certain strains of *S. epidermidis* (Nakatsuji, Chen et al. 2018). Naik *et al*. found that *S. epidermidis* (from the A20 clade) applied to the skin of germ-free mice resulted in the development of protection against specific cutaneous pathogens (Naik, Bouladoux et al. 2012). Other studies have shown that *S. epidermidis* may enhance innate skin immunity and limit pathogen invasion (Naik, Bouladoux et al. 2015). In addition, *S. epidermidis* produces lipoteichoic acid, which can suppress Toll-like receptor (TLR) 3-mediated inflammation and protect mice from *S. aureus* infection (Lai, Cogen et al. 2010, Li, Lei et al. 2013). Moreover, recent evidence suggests that topical *S. epidermidis* application is able to induce a non-classical MHC-I-restricted immune response to not only promote protection against pathogens, but also accelerate wound repair in skin (Linehan, Harrison et al. 2018). The introduction of *S. epidermidis* could also aid in re-establishing skin homeostasis by targeting pathogen-driven dysbiosis.

Translational use of *S. epidermidis* in humans ideally requires controlling the viability and growth of the microbe, particularly in the manipulation, formulation, and lifespan of the therapeutic strain. While genetic manipulation of the strain using classical antibiotic selection markers could achieve this control, these are discouraged by the Food and Drug Administration (FDA) to comprise the final design of an LBP (Dreher-Lesnick, Stibitz et al. 2017). Here, we propose an elegant solution for controlling an *S. epidermidis* strain for therapeutic use by introducing a D-alanine auxotrophy. D-alanine is required for the synthesis of peptidoglycans, which are essential components of the bacterial cell wall. D-alanine is not normally present at significant levels in human tissue, so bacteria must produce it biosynthetically or it must be supplied exogenously. In bacteria, biosynthesis is accomplished through the action of alanine racemase, which interconverts L-alanine and D-alanine, and by D-alanine aminotransferase, which interconverts D-glutamic acid and D-alanine. The metabolic pathway requirement for D-alanine auxotrophy varies in different genetic backgrounds. It was reported that the presence of glutamate racemase (interconverting D-glutamate and L-glutamate) with D-alanine aminotransferase (interconverting D-alanine and D-glutamate) provides a bypass for alanine racemase in *S. aureus* and *Listeria monocytogenes* (Thompson, Bouwer et al. 1998, Moscoso, Garcia et al. 2018). In the present study we found that, in *S. epidermidis*, it was necessary to knock out the D-alanine aminotransferase gene (SE1423) in addition to the two alanine racemase enzymes, *alr1* (SE1674) and *alr2* (SE1079) to fully confer the auxotrophic phenotype.

Here, we describe a strategy for generating a D-alanine auxotrophic strain of *S. epidermidis*. This approach permitted tunable and precise control of growth of the organism. We further characterized its growth in culture and human blood, and its colonization and activity in cultured skin models *in vitro*.

## MATERIALS AND METHODS

### Generation and Confirmation of Auxotrophy

The commensal, non-pathogenic strain of *S. epidermidis* Northern Regional Research Laboratory (NRRL) B-4268 was obtained from the USDA (Agricultural Research Service, NRRL culture collection, Peoria, IL, USA). This strain is also known as ATCC 12228 (American Type Culture Collection [ATCC], Manassas, VA, USA) and PCI 1200 (US FDA). This strain was selected for its low virulence potential because it lacks the *ica* operon implicated in *S. epidermidis-*associated catheter bloodstream infections (Wei, Cao et al. 2006).

The reference genome has been published as the ATCC 12228 genome (Zhang, Ren et al. 2003, MacLea and Trachtenberg 2017). The genome encodes two annotated alanine racemase genes, SE1674 (*alr1*) and SE1079 (*alr2*); both genes were targeted for deletion. The strategy for conferring D-alanine auxotrophy to NRRL B-4268 is presented in **Fig 1A**. Briefly, chromosomal DNA from *S. epidermidis* NRRL B-4268 was isolated and used as a template for PCR. The 5’ flanking region (1.0 Kb, 974 bp) of SE1674 was amplified using forward primer 1674-5F(SalI) and reverse primer 1674-5R. Similarly, the 3’ flanking region (1.0 (998 bp) Kb) of SE1674 was amplified using forward primer 1674-3F and reverse primer 1674-3R (EcoR1). Overlapping PCR was performed using a mixture of the 5’-PCR and 3’-PCR products as templates, and primers 1674-5F (SalI) and 1674-3R (EcoRI) to generate a PCR product 5’-1674-3’ (2.0 Kb).

**FIGURE 1.**
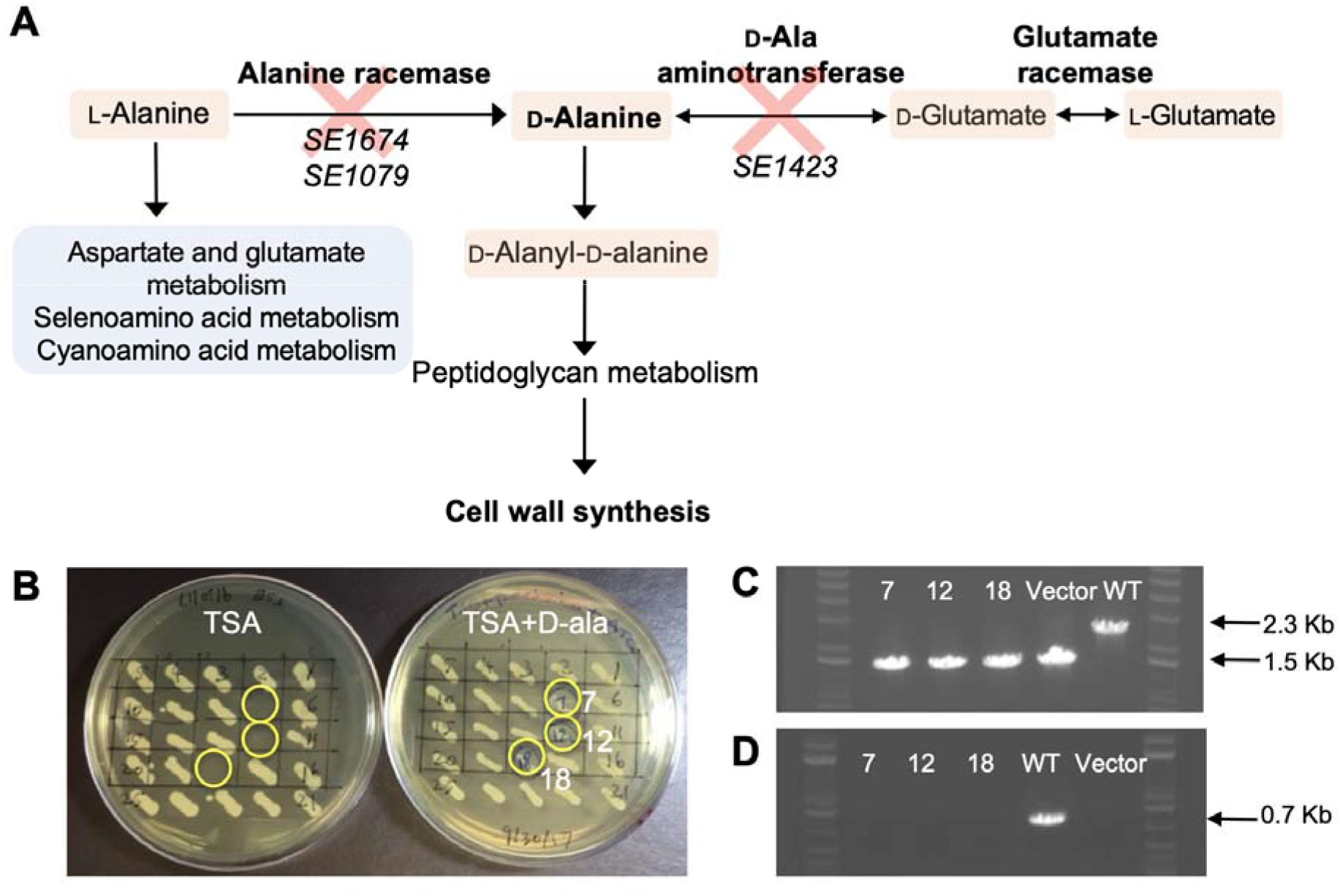
Strategy for D-alanine auxotrophy. (A) Alanine metabolism in *staphylococcal* species and the strategy for constructing a *S. epidermidis* D-alanine auxotroph. (B) The construction of SEΔ*alr*1Δ*alr2*Δ*dat* is described in Methods. Twenty-five candidate clones were patched onto two different plates, and the plates were incubated at 30 °C overnight. Left: TSA plate; Right: TSA + Anhydrotetracycline (2 μg/mL) + D-alanine (40 μg/mL). Three clones (#7, #12 and #18, highlighted in yellow circles) could only grow on TSA supplemented with D-alanine. (C) and (D): Example of PCR confirmation of gene knockout for SE 1423 in SE_ΔΔΔ_ candidates. The DNA from cells patched onto a plate of TSA + Anhydrotetracycline (2 μg/mL) + D-alanine (40 μg/mL) was used as template in PCR reactions. Gel lanes: KO Clone #7; KO Clone #12; KO Clone #18; Wild type SE; SE1423KO plasmid DNA (Vector, as control). For (C) PCR was performed using primers 1423-5F and 1423-3R to distinguish wild type SE1423 locus (PCR product of 2.3 Kb) and SE1423 knockout (PCR product of 1.5 Kb). For (D) PCR was performed using primers 1423-F and 1423-R to detect a PCR product of 0.7 Kb, specific for the wild type SE1423 locus. As expected the PCR product was not generated from the SE1423 knockout plasmid and putative SE1423 knockout *S. epidermidis* clones. Results confirmed successful SE1423 deletion in Clones #7, #12 and #18 and similar PCR analysis was done to confirm deletions of SE1674 and SE1079.

The overlapping PCR product 5’-1674-3’ was digested with SalI and EcoRI and cloned into the temperature-sensitive plasmid pJB38 at the EcoRI-SalI sites (Bose, Fey et al. 2013) and transformed into Top10 *Escherichia coli* (Life Technologies, Inc., Carlsbad, CA, USA) using ampicillin selection (100 μg/mL) to develop knockout (KO) plasmid pJB-1674KO that contains a chloramphenicol (Cm) resistance selection marker. Plasmid pJB-1674KO was purified and transformed into *E. coli* GM2163 (*dam*^−^/*dcm*^−^) using ampicillin selection (100 μg/mL), and then purified and transformed into NRRL B-4268 by electroporation. Transformants were grown at the permissive temperature (30°C) on tryptic soy agar plates (TSA) containing 10 μg/mL Cm. (See **Supplemental Tables S1 and S2** for more information on the strains, plasmids, and primers used.)

For each of the gene knockouts created in this study, the presence of each pJB38 KO plasmid in initial *S. epidermidis* transformants was confirmed by PCR amplification of the flanking region of each gene. Colonies of confirmed pJB38 KO plasmid transformants were restreaked onto fresh TSA + Cm (10 μg/mL) + D-alanine (40 μg/mL) and plates were incubated at 43°C for 24h to select for colonies with plasmid integrated into the chromosome via single-crossover homologous recombination. Isolated colonies were streaked again for purification at 43°C and then inoculated into 50 mL tryptic-soy broth (TSB) + D-alanine (40 μg/mL), without Cm, in a 250 mL baffled shake flask and grown at 30°C for 24h to permit the excision of plasmid sequences from the chromosome by homologous recombination. The culture was passaged six times by transferring an aliquot of 0.5 mL to a flask containing 50 mL fresh medium, and growing at 30°C for 24 h. The final culture was plated on TSA + anhydrotetracycline (2 μg/mL; to induce plasmid counter-selection) + D-alanine (40 μg/mL). After two days of incubation at 30°C, approximately 100-200 colonies were formed on plates inoculated with 100 μL of culture at 10^−5^ dilution. Following integration of the KO plasmid and removal of the plasmid backbone containing the antibiotic selection marker, an SE1674 knockout strain (SEΔ*alr1*) was isolated and confirmed by PCR amplification using primers flanking the gene. Using the same gene knockout method described above, we proceeded to knockout SE1079 in this strain, leading to successful generation of double knockout strains (SEΔ*alr1*Δ*alr2*). More information about the plasmids and PCR primers used in this study can be found in **Supplemental Tables S1 and S2**, respectively.

When SEΔ*alr1*Δ*alr2* was found to grow normally in the absence of D-alanine supplementation to the media, we hypothesized that D-alanine aminotransferase (*dat*, SE1423) could provide a bypass for alanine racemase in *S. epidermidis*, as was suggested for S. epidermitdis D-alanine metabolism pathway by KEGG (https://www.genome.jp/dbget-bin/www_bget?sep00473), and reported in *S. aureus* and *Listeria monocytogenes* (Thompson, Bouwer et al. 1998, Moscoso, Garcia et al. 2018). Employing the same gene knockout strategy and method mentioned above, a triple knockout *S. epidermidis* strain (SEΔ*alr1*Δ*alr2*Δ*dat*, or SE_ΔΔΔ_) was constructed, which could not grow in the absence of exogenously supplied D-alanine.

### Measurement of Effect of D-Alanine Levels on SE_ΔΔΔ_ Growth

The effect of D-alanine concentrations ranging from 0 μg/mL to 200 μg/mL (0% to 0.02%) on the short-term growth curves of SE_ΔΔΔ_ was investigated. Five mL of Vegitone medium (Sigma Aldrich, catalogue no. 41960) containing 0.004% (40 μg/mL) of D-alanine (ACROS Organics, 99+%) was inoculated with an isolated SE_ΔΔΔ_ colony. Cultures were incubated overnight at 37°C, shaking at 250 revolutions per minute (RPM) on an orbital shaker.

Growth rates were measured at 37°C for up to 7 h. Fifty mL of Vegitone containing the prescribed amount of D-alanine was inoculated with 1 mL of the overnight culture to give a final optical density (OD) at 600nm of 0.1 relative units (ru). Flasks were incubated on orbital shaker at 250 RPM. Every ~60 min, a sample was taken and the OD_600_ recorded. Growth curves were conducted for 7h and were fitted using a standard logistic growth equation (Yin, Goudriaan et al. 2003).

### Measurement of Effect of Temperature and pH on SE_ΔΔΔ_ Growth

The effect of temperature and media pH on SE_ΔΔΔ_ growth were examined, using 100 μg/mL D-alanine in all groups. Growth curves were measured at room temperature (~25°C) and 37°C for up to 40h. Fifty mL of Vegitone containing D-alanine was inoculated with 1.0 mL of the overnight culture. Growth curves were also evaluated at three different pH values (4, 5, 6, and 7) at 37°C for 40 h.

### Antibiotic susceptibility testing of *S. epidermidis* NRRL B-4268 and SE_ΔΔΔ_

Antibiotic susceptibility testing was conducted at IHMA Europe Sàrl, Monthey, Switzerland using Clinical and Laboratory Standards Institute (CLSI) methodology, with the exception that 100 mg/L D-alanine was added to testing with SE_ΔΔΔ_ (Weinstein and Lewis 2020).

### Determination of the Frequency of Spontaneous Phenotypic Reversion of Auxotrophy of SE_ΔΔΔ_

The D-alanine deficient media used to isolate revertants was Vegitone agar plates, prepared in 150 mm petri dishes. Rifampin stocks were prepared in methanol at a final concentration of 4 μg/mL. As appropriate, Vegitone agar plates were prepared with either 100 μg/mL of D-alanine, or 8 ng/mL of rifampicin and 100 μg/mL of D-alanine. Eight milliliters of the overnight culture were spun down at 10,000 rpm for 10 min. The pellets were resuspended in 1200 μL of Vegitone broth and 150 μL of the suspension was spread onto the above 150 mm petri dishes. All plates were incubated at 37°C for ~24h. The experiment was carried out in quadruplicate. The remaining inoculum suspension was serially diluted 1:10 in Vegitone broth and plated onto Vegitone agar plates supplemented with 100 μg/ml of D-alanine and incubated at 37°C for 24h to quantify colony forming units (CFUs) in the inoculum. The spontaneous resistance frequency was determined by dividing the number of colonies grow in in the absence of D-alanine by the total number of CFUs plated.

### Determination of Growth of SE_ΔΔΔ_ in Blood

TSB supplemented with 100 μg/mL of D-alanine was inoculated with SE_ΔΔΔ_ and incubated overnight at 37°C. The overnight culture was further diluted in fresh TSB + D-alanine and allowed to grow until it reached mid-log phase, which corresponded to an absorbance of ~0.7, measured by OD_600_. Virus-free, defibrinated pooled whole human blood from three healthy donors was purchased from BioIVT (Westbury, NY). The SE_ΔΔΔ_ culture was concentrated to 1×10^9^ CFU/mL. Serial ten-fold dilutions ranging from 1×10^9^ to 1×10^1^ were made in TSB ± D-alanine, then diluted again ten-fold into a volume of 100 μL blood in a 96-well microtiter plate. All blood cultures were incubated in a tissue culture incubator at 37°C, 5% CO_2_ for 24h. The following day, cultures were serially diluted and plated on TSA with 100 μg/mL of D-alanine, grown overnight, and scored for the number of colonies. The experiment was carried out in duplicate. The remaining inoculum suspension was serially diluted in TSB and plated onto TSA supplemented with 100 μg/mL of D-alanine to quantify CFUs in the inoculum. All plates were incubated at 37°C for 24h.

### Growth of SE_ΔΔΔ_ on Reconstructed Human Epidermis

Fully differentiated antibiotic-free reconstructed human epidermis (RHE) culture inserts from Mattek (EpiDerm, Ashland, MA, USA) were used to evaluate bacterial colonization. RHE models were reactivated by placing cell inserts into 6-well plates containing 1 mL of EpiDerm maintenance media from Mattek. The inserts were then incubated for 18h at 37°C in 5% CO_2_. Following incubation, the inserts were transferred to 12-well hang tops with 5 mL of Epiderm maintenance media in each well, and the tissues were incubated for an additional 30 min at 37°C in 5% CO_2_. When applicable, 100 μg/mL of D-alanine was also added to the RHE media. SE_ΔΔΔ_ was inoculated into 25 mL of TSB containing 100 μg/ml of D-alanine. The culture was grown at 37°C overnight while shaking. After 18h, fresh TSB supplemented with 100 μg/mL of D-alanine was inoculated with the overnight culture of SE_ΔΔΔ_, at a starting OD_600_ of 0.1. Cultures were grown at 37°C to log phase (OD_600_ 0.5-0.7) and pelleted by centrifugation at 15,000 rpm for 2 min, then resuspended in Dulbecco’s phosphate buffered saline (DPBS) to a final OD_600_ of 1.0. The bacterial pellet was suspended in 1 mL of DPBS and pelleted at 15,000 rpm for 2 min. This process was repeated two more times, to ensure residual D-alanine was removed from the bacterial suspension. The bacterial cells were next diluted down to 0.01 OD_600_ in DPBS. For cells receiving D-alanine, 100 μg/mL of D-alanine was added to the bacterial suspension prior to RHE inoculation. To inoculate the RHE inserts, 10 μL of the prepared bacterial suspension was applied to the center of the RHE ensuring that the liquid droplet remained centered when possible. The RHE inserts were then incubated at 37°C in 5% CO_2_ for 4h, 24h, 48h, or 72h as indicated. At each time point bacterial colonization was assessed by taking a 4 mm punch biopsy from the center of RHE. Biopsies were washed by submerging the tissue in 1 mL DPBS to remove unadhered cells. In order to release the colonized bacterial cells from the RHE, the biopsies were transferred to 1 mL of DPBS, followed by vortexing for 2 min at full speed. Colonization of SE_ΔΔΔ_ was quantified by dilution plating and counting colony forming units (CFU) on SASelect plates supplemented with 100 μg/mL D-alanine (BioRad, Hercules, CA). All experiments were done in triplicate.

### Biofilm Formation Assay with SE_ΔΔΔ_

The biofilm assay was adapted from Mack et al, 1992 and Neopane et al, 2018 (Mack, Siemssen et al. 1992, Neopane, Nepal et al. 2018). TSB was inoculated with *S. epidermidis* strains SE_ΔΔΔ_ and the biofilm-forming strain *S. epidermidis* 1457 (Galac, Stam et al. 2017) and incubated overnight at 37°C. The culture was then diluted to 1:200 in fresh TSB/0.5% glucose and 200 μL dispensed into a flat bottom 96-well tissue culture plate (Thermo Fisher Scientific, Waltham, MA). The plate was incubated at 37°C for 24h. After incubation, the microtiter plate was washed twice with 1x DPBS to remove loosely attached bacterial and planktonic cells. The biofilms were fixed by incubating in 200 μL of 99% methanol for ten min. The methanol was decanted and wells were allowed to air dry for ten min. Once dry, wells were stained with 0.1% crystal violet for five min. Excess crystal violet was washed twice with deionized water. Wells were air dried, then stained biofilms were removed by dissolving in 200 μL of 30% glacial acetic acid. The amount of biofilm was quantified by measuring absorbance of the plate at 570 nm.

### Effect of SE_ΔΔΔ_ on Antimicrobial Peptide (AMP) Production in Cultured Human Skin

The effect of SE_ΔΔΔ_ on AMP production in cultured human skin was performed at StratiCELL (Les Isnes, Belgium) using RHE inserts. RHE inserts were cultured in triplicates at the air-liquid interface for 14 days in gentamicin-free culture medium containing D-alanine (100 μg/mL) in 95% humidity at 37°C with 5% CO_2_ to fully differentiate. A single topical treatment with SE (10 μL at a cell density of 1×10^7^ CFU/mL in PBS + 100 μg/mL D-alanine) was allowed to colonize for 48h on fully differentiated epidermis. Control samples were treated with 10 μL of the vehicle solution (PBS + D-alanine 100 μg/mL). During the treatment, the tissue culture medium feeding the RHE was also supplemented with 100 μg/mL D-alanine. RHE inserts were kept at the air-liquid interface in a humid atmosphere at 37°C with 5% CO_2_.

#### Histological Analysis of RHE

At the end of the treatment, three tissues per condition were fixed in 4% formaldehyde, dehydrated and paraffin embedded. Sections of 6 μm of epidermis were stained with eosin and hematoxylin. Slides were mounted with specific medium and examined with a Leica DM2000 photomicroscope coupled to a digital camera (Zeiss).

#### Analysis of AMP Expression in RHE

To quantify gene expression of human AMPs, total mRNA was extracted using the Qiagen RNeasy kit (Hilden, Germany). Tissues were washed in PBS, removed from their inserts and immersed directly in the lysis buffer. Extraction of mRNA was performed according to the supplier’s recommendations. The collected mRNA was stored at −80°C. Reverse transcription was performed with the high capacity mRNA-to-cDNA kit (Applied Biosystems, Foster City, CA) from 1 μg of total mRNA according to the manufacturer's instructions.

The target sequences of the genes of interest, S100 Calcium-binding protein A7 (S100A7) and human *β*-defensin 2 (DEFB4A) were amplified by using TAQMAN Gene Expression Assays (Applied Biosystems, Foster City, CA). TaqMan probes were grafted with a fluorophore (FAM) at their 5’ end and with a fluorescence quencher in 3’. PCR reactions were performedwith the Quantstudio7 Real-Time PCR System (Applied Biosystems, Foster City, CA). In order to normalize the results, a housekeeping gene (β2-microglobulin; B2M) was used. The thermal cycles were programmed with one first denaturation step at 95°C for 20 s. The amplification protocol was followed with 40 cycles (1s at 95°C and 20s at 60°C). Threshold cycles (Ct) were obtained for each gene. Data were analyzed by using the RQ application available on Applied Biosystems website (and designed to perform relative quantification of gene expression using the comparative Ct (ΔΔ Ct) method (Livak and Schmittgen 2001, Pfaffl 2001). Independent, two-sided T-tests were used to test gene expression between treated and untreated RHE.

## RESULTS

### Construction of D-alanine auxotrophic strain SEΔ*alr1*Δ*alr2*Δ*dat* (SE_ΔΔΔ_)

Knocking out of both alanine racemase genes, *alr1* and *alr2*, in *S. epidermidis* NRRL B-4268 (SEΔ*alr1*Δ*alr2*) did not produce an auxotropic phenotype for D-alanine, in contrast to similar genetic knockouts in other bacteria such as *Bacillus subtilis* (Dul and Young 1973). This suggested that there was another metabolic pathway that could result in the production of D-alanine. The *dat* gene, encoding D-alanine aminotransferase, was identified as a candidate gene to knockout in order to eliminate potential interconversion of D-glutamate to D-alanine (**Fig. 1A**). The resulting “triple knockout” strain, SEΔ*alr1*Δ*alr2*Δ*dat* (SE_ΔΔΔ_), was successfully generated and displayed the desired D-alanine auxotrophy. All candidate clones (#7, #12 and #18) grew well on TSA medium supplemented with D-alanine and failed to grow in the absence of supplementation (Fig. 1B). All three chromosomal deletions were similarly confirmed by PCR in each of the three candidate clones, #7, #12 and #18, as illustrated in the example for SE1423 shown in **Fig. 1 C** and **1D**.

### Effect of D-alanine Concentration in Medium on the Growth of SE_ΔΔΔ_ in Broth Cultures

The kinetics of SE_ΔΔΔ_ growth in Vegitone broth were evaluated at 37°C with increasing concentrations of D-alanine. (**Fig. 2A**). After ~7h, growth approached stationary phase, independently of the D-alanine concentration, such that generation times could be estimated using a standard logistic equation. From this, the estimated generation time was ~31-45 min. No growth of the SE_ΔΔΔ_ was measured in the absence of D-alanine after 7h, in agreement with the auxotrophic nature of the bacteria. There was no correlation between generation time and D-alanine concentration. The growth of SE_ΔΔΔ_ was very sensitive to the D-alanine concentration as indicated by a steep hillslope (~11) of the concentration-growth curve (not shown). The D-alanine concentration needed for the half-maximal response (EC_50_) was 54 μg/mL (0.005%) (**Fig. 2B**). As there was no difference in the growth curves between 80 and 200 μg/mL of D-alanine, the standard concentration of D-alanine for routine growth of the auxotroph was set at 100 μg/mL.

**FIGURE 2.**
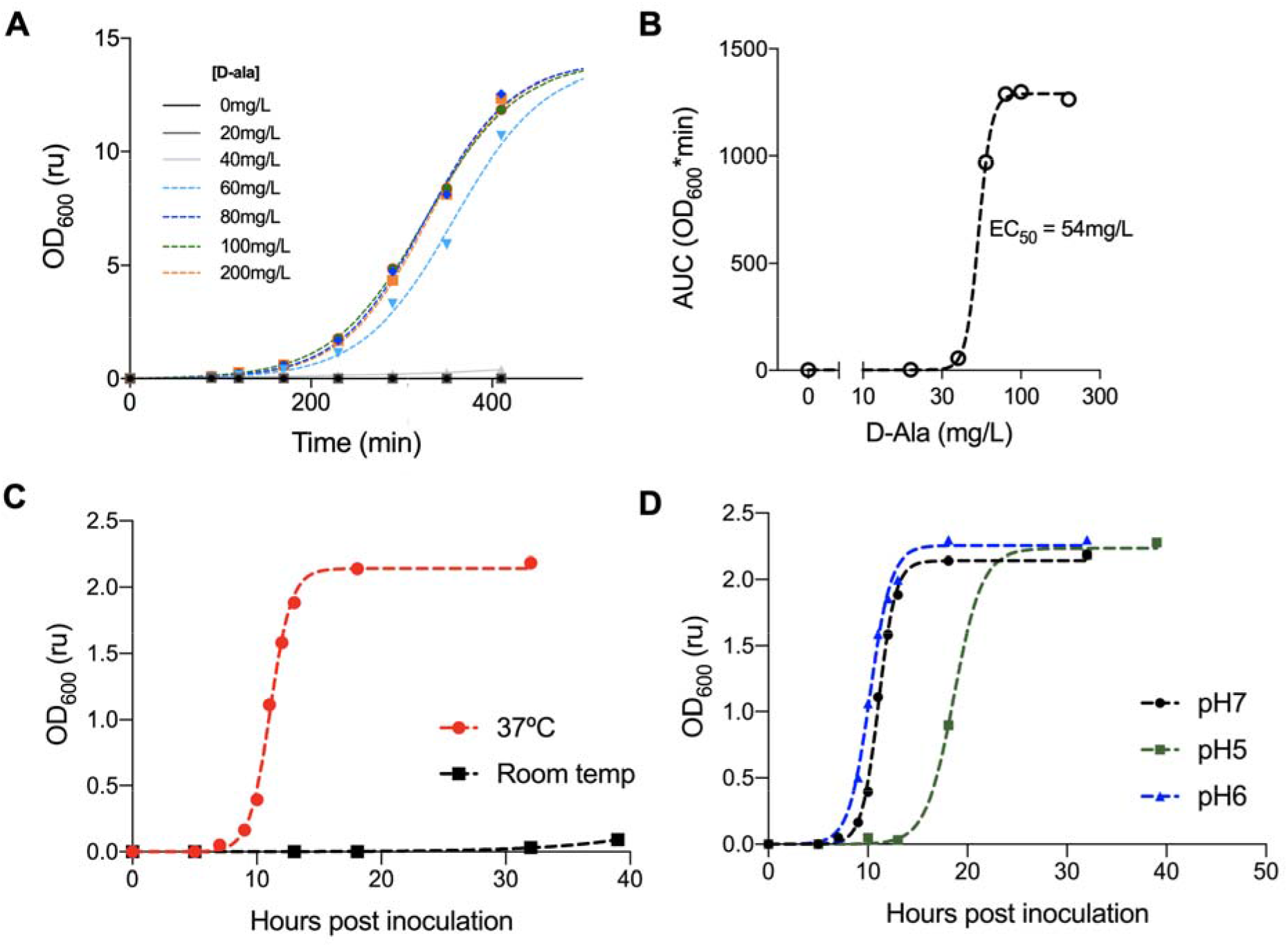
Characterization of SE_ΔΔΔ_ growth *in vitro*. (A) Graph showing the effect of D-alanine concentration (from 0 to 20 μg/mL) on the change in OD at 600nm over time of incubation at 37°C for SE_ΔΔΔ_. Each data point is the mean ± SD of three independent replicates. Dashed lines are the results from the logistic fit. (B) EC_50_ curve of D-alanine concentration on AUC. (C) Graph showing the effect of temperature on the growth kinetic of SE_ΔΔΔ_. Growth curves were generated at room temperature and at 37°C using a D-alanine concentration of 100 μg/mL. One culture was placed into the incubator shaker (37°C; 250 rpm) and the second culture at room temperature in a shaker (250 rpm). (D) Graph showing the effect of pH on the growth kinetic of SE_ΔΔΔ_. Growth curves, at 37°C, were generated from Vegitone media that had been adjusted to pH 5, 6, or the standard pH 7.0. D-alanine was then added to obtain a final concentration of 100 μg/mL.

### Effect of Temperature and pH on the Growth of SE_ΔΔΔ_ in Broth Cultures

At room temperature, SE_ΔΔΔ_ showed little to no growth as monitored by OD_600_ **(Fig. 2C)**, though the culture density increased by ~2 log_10_ CFUs over a 30h time period (not shown). The comparator culture grown at 37°C had a log_10_ CFU increase of ~6 (not shown) and reached stationary phase within ~4h from the end of lag phase (**Fig. 2C**).

Skin surface pH ranges from ~ 4-6, as compared to below the skin surface (Ali and Yosipovitch 2013), therefore growth at lower pH in broth cultures was evaluated. SE_ΔΔΔ_ cultures grown at 37°C in standard Vegitone broth pH (7.0) and in broth adjusted to pH 6.0 each increased by ~6 log_10_ CFUs (not shown) and reached stationary phase within 4h from end of lag phase, showing generation times of 45.5 min and 35.5 min, respectively (**Fig. 2D**). Growth rate at pH 5.0 was somewhat slower (generation time of 65 min) and showed a more pronounced lag phase yet grew to about the same final cell density as with media at pH 6 and 7 **(Fig. 2D)**. SE_ΔΔΔ_ showed no growth at pH 4 (data not shown).

### Spontaneous Reversion of D-alanine Auxotrophic Phenotype

In order to determine the frequency of reversion from D-alanine auxotrophy to prototrophy of SE_ΔΔΔ_, concentrated cell suspensions were plated on Vegitone agar without D-alanine supplement and compared to plates with D-alanine. The result indicates that no phenotypic revertants developed in the absence of D-alanine, with an estimated frequency of reversion less than 4.8 × 10^−11^ revertants/CFU plated. In the same strain, the frequency of spontaneous rifampicin resistance emergence was 6 × 10^−8^ mutants/CFU plated, demonstrating that the conditions of this experiment were able to select for mutations in strain SE_ΔΔΔ_ (**Supplementary Fig. S1**).

### Growth in Human Blood

To test the ability of SE_ΔΔΔ_ to grow inappropriately should it enter the bloodstream, growth was assessed for 24h in defibrinated human blood. SE_ΔΔΔ_ inoculated in human blood without D-alanine showed no growth after an overnight incubation for inoculum less than 1.4 × 10^8^ CFU/mL, but human blood supplemented with D-alanine was capable of supporting growth at all levels of inocula tested (except the lowest inoculum concentration of 1 × 10^0^) (**Supplementary Table S3**). The CFU counts of blood cultures inoculated with 1.4 × 10^8^ CFU/mL remained constant, with no further growth or loss of viability under the conditions of this assay.

### Antibiotic Susceptibility of SE_ΔΔΔ_

The antibiotic susceptibility profiles of NRRL B-4268 and SE_ΔΔΔ_ were determined using CLSI methodology. **Table 1** shows that the SE_ΔΔΔ_ strain has antibiotic susceptibility similar to the parent wild type *S. epidermidis* NRRL B-4268, and results were consistent with a prior literature report (Zhang, Ren et al. 2003).

**Table 1:** Antibiotic Susceptibility of SE_ΔΔΔ_ Versus Parent *S. epidermidis*.

| Antibiotic | MIC (μg/mL) |  | CLSI S/I/R<sup>a</sup> |
| --- | --- | --- | --- |
|  | SE<sub>ΔΔΔ</sub> | SE NRRL B-4268 |  |
| Ampicillin | 4 | 4 | N/A |
| Bacitracin | 16 | 16 | N/A |
| Cefalexin | 2 | 2 | N/A |
| Ceftaroline | 0.12 | 0.12 | S |
| Chloramphenicol | 2 | 2 | S |
| Clindamycin | 0.06 | 0.06 | S |
| Daptomycin | 0.5 | 0.5 | S |
| Doxycycline | 4 | 4 | S |
| Erythromycin | ≤0.12 | ≤0.12 | S |
| Fosfomycin | 4 | 4 | N/A |
| Fusidic acid | ≤0.03 | ≤0.03 | N/A |
| Gentamicin | ≤0.06 | ≤0.06 | S |
| Levofloxacin | 0.12 | 0.12 | S |
| Linezolid | ≤0.5 | ≤0.5 | S |
| Mupirocin | 0.12 | 0.12 | N/A |
| Oritavancin | 0.03 | 0.06 | S |
| Oxacillin | 0.12 | 0.12 | S |
| Quinupristin/Dalfopristin | ≤0.12 | ≤0.12 | S |
| Rifampicin | ≤0.002 | ≤0.002 | S |
| Tetracycline | >32 | >32 | R |
| Tigecycline | 0.25 | 0.12 - 0.25 | S |
| Trimethoprim/Sulfamethoxazole | ≤0.5/9.5 | ≤0.5/9.5 | S |
| Vancomycin | 1 | 1 | S |

\* s – sensitive ; R - resistant

### SE_ΔΔΔ_ Colonization of a Human Skin Model *in vitro*

To determine if D-alanine supplementation is necessary for SE_ΔΔΔ_ colonization and survival on the RHE, 10^5^ CFU of SE_ΔΔΔ_ were inoculated onto the RHE surface either with or without D-alanine supplementation. No colonization occurred without D-alanine present, indicating that D-alanine must be supplied for growth on skin (**Fig. 3**). Furthermore, these data show that SE_ΔΔΔ_ colonized the RHE within 4h and persisted for up to 72h following a single application with D-alanine supplementation. Interestingly, as early as 4h post inoculation, in the absence of D-alanine, no bacteria were recovered from the RHE when homogenates were plated on SAselect plates supplemented with D-alanine.

**FIGURE 3.**
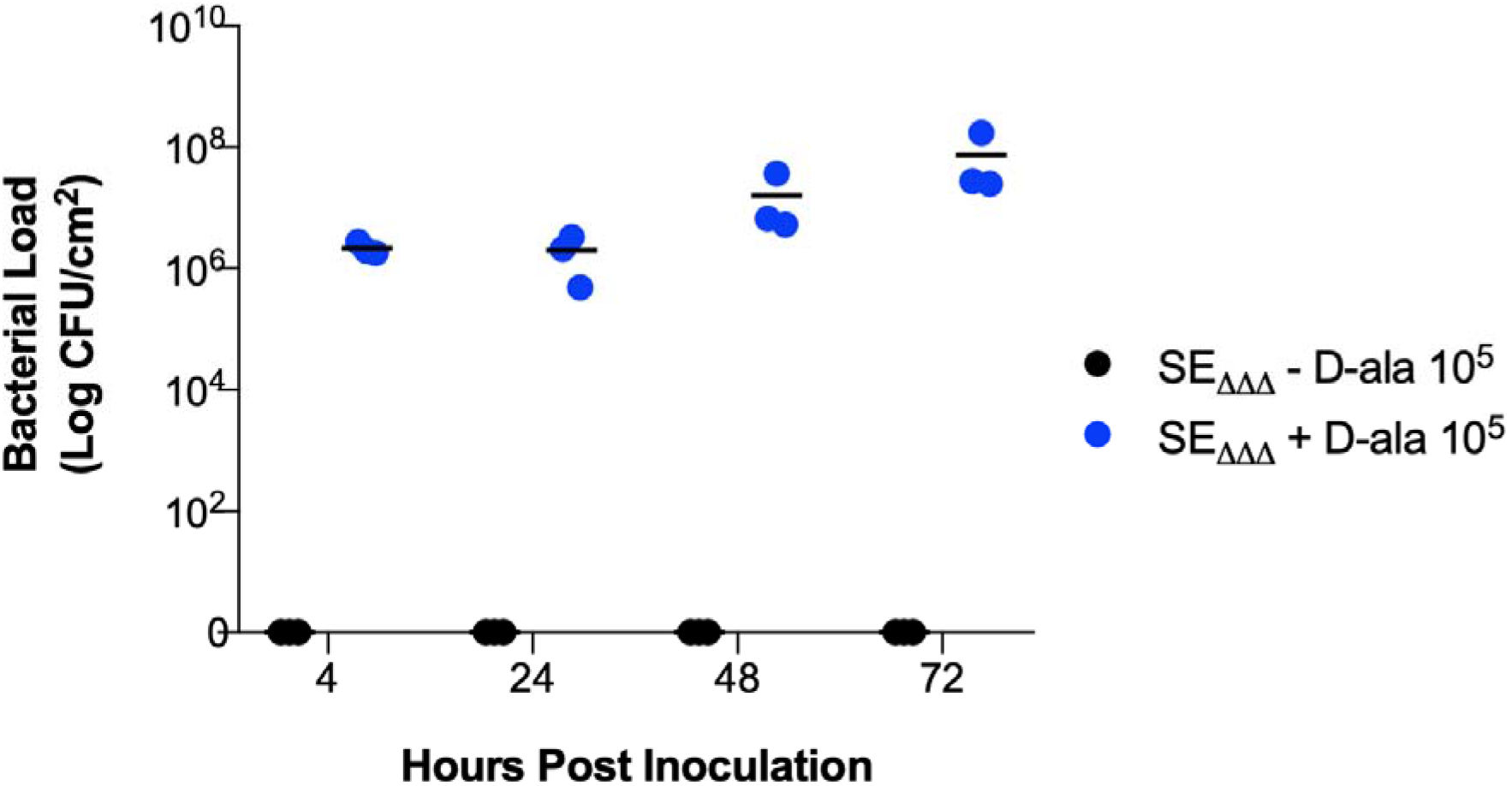
SE_ΔΔΔ_ Growth on RHE with and without D-Alanine Supplementation. RHE inserts were colonized with SE_ΔΔΔ_ at 37ºC, 5% CO_2_. When present, D-alanine, 100 μg/mL, was added to the media feeding the RHE insert. Colonized models were harvested at each time point by uniform skin punches. Skin punch samples were rinsed in DPBS to remove non-adherent cells, vortexed, and assayed for cell density by serial dilution and plating on SAselect plates supplemented with 100 μg/mL of D-alanine. Each condition was tested in triplicate.

### Biofilm formation on plastic

The ability of SE_ΔΔΔ_ to form biofilm on polystyrene plastic was evaluated in a standard crystal violet assay in a 96-well format. Cultures were grown in TSB/0.5% D-glucose, with and without D-alanine, and biofilm formation was detected and quantified using crystal violet. **Fig. 4** shows that the crystal violet staining for SE_ΔΔΔ_with or without D-alanine present was indistinguishable from the blank and TSB controls, indicating no biofilms formed on the plastic under the conditions of this assay. Wells containing the positive control strain, SE1457, showed strong crystal violet staining, consistent with this strain’s known *ica-*positive genotype and biofilm-forming phenotype.

**FIGURE 4.**
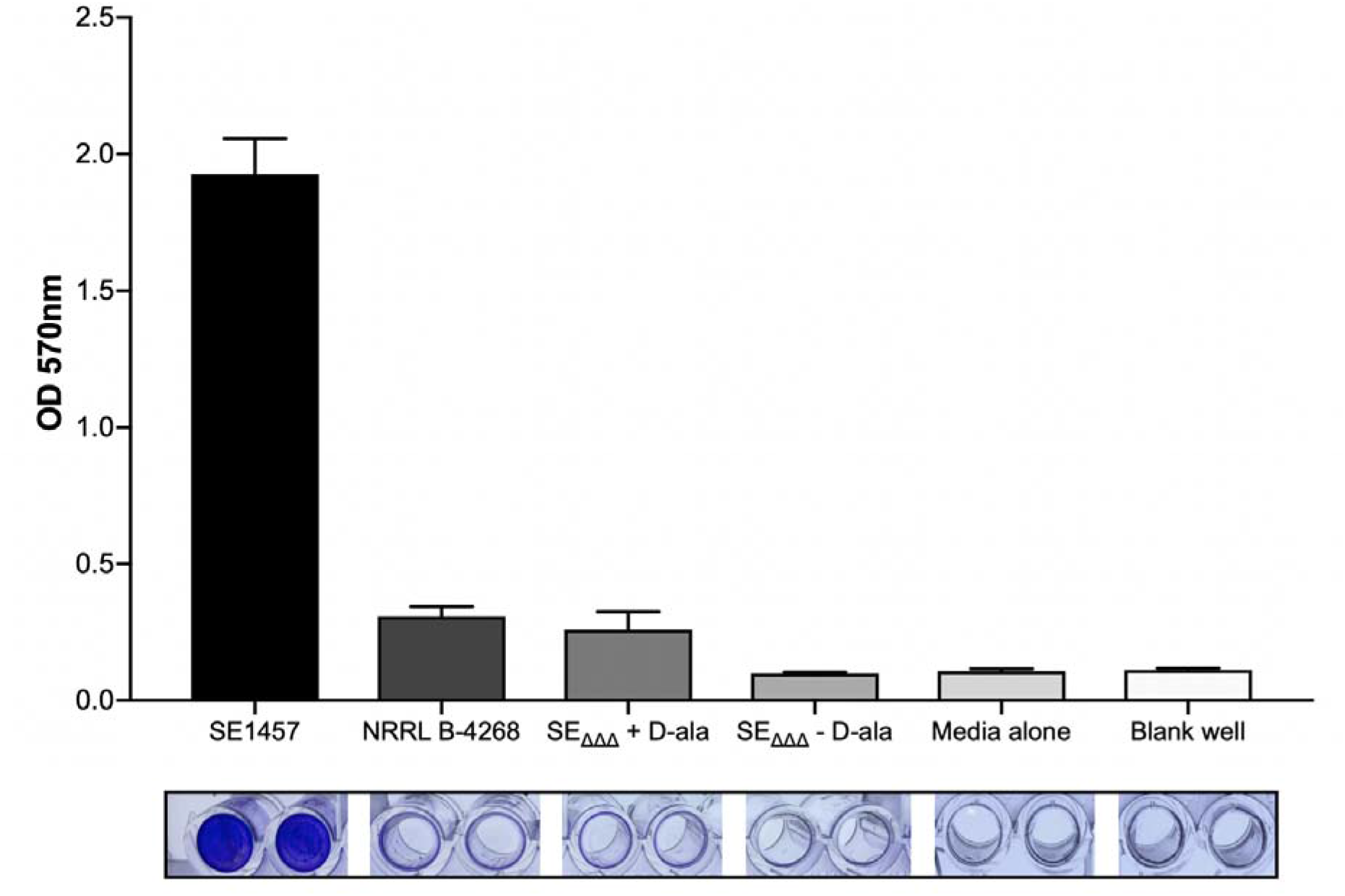
The production of biofilm from *S. epidermidis in vitro*. The bottom photograph panel shows the accumulation of biofilm on plastic wells within 24h at 37°C, visualized by crystal violet staining. Biofilm formation was further quantified by dissolving the crystal violet in acetic acid and reading absorbance at 570 nm. *S. epidermidis* strain 1457 was the positive control, and the blank and TSB media-only wells served as negative controls for background staining. All data show mean ± SD of 16 replicate microtiter wells.

### Effect of SE_ΔΔΔ_ on human AMP expression

SE_ΔΔΔ_ was also evaluated for its ability to trigger a host-commensal communication. SE_ΔΔΔ_ was seeded on RHE inserts and mRNA levels for specific host-commensal signals were measured by qPCR. **Fig. 5** shows the change in expression levels of two epithelial AMPs, S100 calcium-binding protein A7 (S100A7) and hβD-2. In this sample (n = 3), the average expression of both AMPs was heightened in the presence of SE_ΔΔΔ_ but only hβD-2 reached statistical significance (229% increase; 95% confidence interval [CI]: 1.14, 4.59; p = 0.047). There was a non-statistically significant increase in expression of S100A7 (fold increase: 2.07; 95% CI: 0.12, 36.0). No deleterious effects of the SE_ΔΔΔ_ colonization on the structure of the RHE was observed histologically.

**FIGURE 5.**
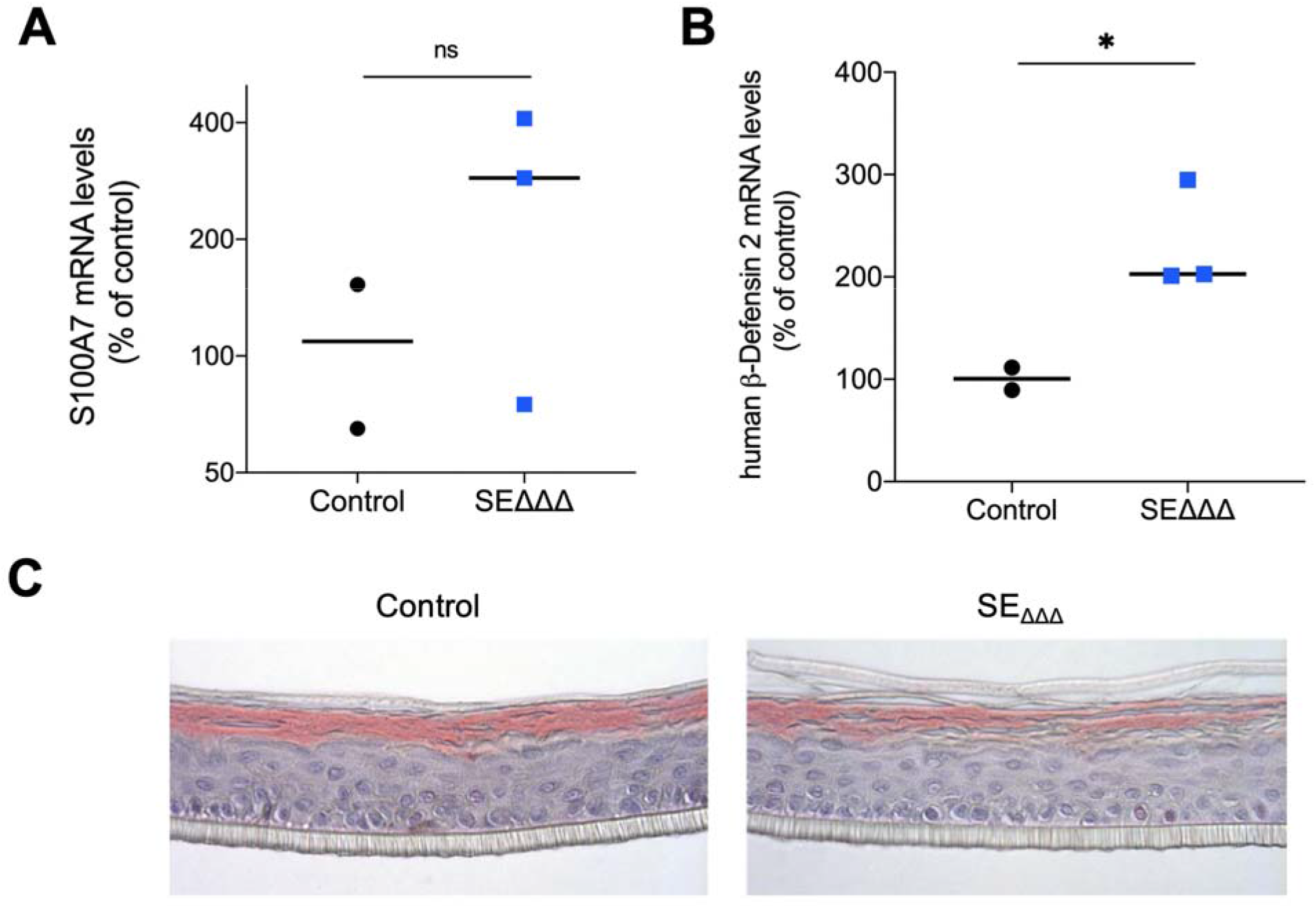
Expression of human AMPs following SE_ΔΔΔ_ treatment of RHE. Panels (A) and (B) show plots of the changes in mRNA levels for two epithelial AMPs (S100A7 and hβD-2). hβD-2 was significantly elevated in the presence of SE_ΔΔΔ_ (* *p*<0.05, T-test). Panel (C) shows hematoxylin and eosin staining of transverse section of RHE inserts with or without treatment with SE_ΔΔΔ_.

## DISCUSSION

A strategy is presented for the construction of a *S. epidermidis* strain, SE_ΔΔΔ,_ with a non-antibiotic-associated conditional growth phenotype. SE_ΔΔΔ,_ exhibits key features that support its potential as a skin LBP: its growth is tunable with D-alanine, its D-alanine auxotrophy has a low propensity to phenotypically revert, it colonizes human skin equivalents, it doesn’t grow in pooled human blood, it doesn’t form biofilms on plastic, and it increases expression of an important host AMP, hβD-2. Our strategy involved deleting the ability to synthesize D-alanine, which is required for the synthesis of bacterial cell peptidoglycan. Here we show that, as expected, disruption of peptidoglycan through D-alanine starvation is bactericidal in culture and on skin, *in vitro*. In contrast to reports for *B. subtilis*, in order to develop D-alanine auxotrophy in SE, not only the two alanine racemase genes (*alr1, alr2*) but also the D-alanine aminotransferase gene *dat* had to be knocked out. Evidently, the combination of glutamate racemase and D-alanine aminotransferase provides a viable bypass to alanine racemase, as reported in MRSA132 (Moscoso, García et al. 2018) and *Listeria monocytogenes* (Thompson, Bouwer et al. 1998). Others have reported on other auxotrophy strategies for attenuating growth in *S. aureus*, including proline (Li, Sun et al. 2010), menadione (Lannergård, von Eiff et al. 2008), and NK41 (Barbagelata, Alvarez et al. 2011).

Several findings from these studies support the idea that attenuation by D-alanine auxotrophy may promote the safety of SE_ΔΔΔ_ for use as a skin LBP. The very low spontaneous reversion rate of D-alanine auxotrophy suggests that very large doses of SE_ΔΔΔ_ may be applied to skin without selecting for a prototrophic revertant. Further, results also support that the D-alanine auxotrophy of SE_ΔΔΔ_ may mitigate the risk of bacterial dissemination into the bloodstream. Biofilm forming SE are often associated with opportunistic nosocomial infections due to the ability to colonize biomaterials of indwelling medical devices (Conlan, Mijares et al. 2012, Kord, Ardebili et al. 2018). Therefore, biofilm formation is an undesirable property of an LBP. No biofilm formation was seen with SE_ΔΔΔ_, with and without D-alanine, suggesting that it may have a lower propensity to form biofilm in a clinical setting. Another important observation is that the inability to form biofilm *in vitro* did not affect the ability of SE_ΔΔΔ_ to robustly colonize and grow on RHE.

*S. epidermidis* belongs to the group of coagulase-negative staphylococci (CoNS), which is distinguished from coagulase-positive staphylococci such as *S. aureus* by lacking the enzyme coagulase. A large arsenal of tools are available to *S. epidermidis* to control the colonization of other pathogenic microorganisms, including proteases that degrade proteins in *S. aureus* biofilm, stimulation of AMP secretion from keratinocytes, or secretion of lantibiotics targeting *S. aureus* (Sahl 1998, Dubin, Chmiel et al. 2001, Moon, Banbula et al. 2001, Bastos, Ceotto et al. 2009, Chen, Krishnan et al. 2013, Sugimoto, Iwamoto et al. 2013, Vandecandelaere, Depuydt et al. 2014). In addition, *S. epidermidis* produces lipoteichoic acid, which can suppress TLR3-mediated inflammation and protect mice from *S. aureus* infection (Lai, Di Nardo et al. 2009). Specific novel lantibiotics secreted by *S. epidermidis* and *Staphylococcus hominis* were recently shown to control *S. aureus* infection by synergizing with the human keratinocyte-secreting cathelicidin-related AMP LL◻37 (Nakatsuji, Chen et al. 2017). This is a further demonstration of the important and beneficial interplay between commensals and the host. Interestingly, these lantibiotics-secreting strains were significantly underrepresented in the microbiome of atopic dermatitis patients and exogenous application of these lantibiotics-secreting commensal resulted in a reduction in *S. aureus* infection in a small group of atopic dermatitis patients (Nakatsuji, Chen et al. 2017). Introduction of D-alanine auxotrophy into other *S. epidermidis* strain backgrounds may be a way of broadly controlling isolates selected for their novel therapeutic properties.

Skin innate immunity is the first line of defense after the physical skin barrier. Keratinocytes are the main skin cell population involved (Pivarcsi, Kemény et al. 2004, Pasparakis, Haase et al. 2014, Bitschar, Wolz et al. 2017); they express a variety of TLRs, which are primary sensors of innate immunity (Lee, Blaber et al. 2011, Lee and Blaber 2013). The innate immune system can recognize pathogens and trigger the host response to eliminate them via the release cytokines (e.g., IL1-α and IL1-β) and AMPs (Pasparakis, Haase et al. 2014, Simanski, Glaser et al. 2014, Wang, Tai et al. 2019, Wei, Wiggins et al. 2019). AMPs are secreted by keratinocytes, sebocytes, T-cells and mast cells and can directly attack pathogens (e.g., bacteria, enveloped viruses, fungi) (Brogden 2005, Schittek, Paulmann et al. 2008, Yamasaki and Gallo 2008, Clausen and Agner 2016). AMPs can be grouped into five separate classes: defensins, dermcidin, cathelicidins, S100 proteins and ribonucleases (Yamasaki and Gallo 2008, Wanke, Steffen et al. 2011, Niyonsaba, Kiatsurayanon et al. 2017). Some are constitutively expressed while others can be induced following a skin insult. AMPs are highly cationic and thus interact with negatively charged membranes components [e.g. lipopolysaccharide (LPS), peptidoglycans, outer membrane protein I (OprI)] of skin pathogens, essentially forming holes in the cell membrane (Brogden 2005). AMPs have also been shown to exert immunomodulatory effect, interacting directly with receptors on immune cells to promote chemotaxis, differentiation and cytokine production (Conlan, Mijares et al. 2012). The most abundant AMPs in human skin are human defensins, the cathelicidin LL-37, and RNase 7. Several studies have shown that commensal organisms such as *S. epidermidis* can directly induce expression of hβD-2 (*hDEFB4A* gene) while presenting a relative tolerance to it, enabling such organisms to survive on the skin surface and modulate defensin expression when the stratum corneum barrier is disrupted (Dinulos, Mentele et al. 2003). Therefore, commensal bacteria such as *S. epidermidis* are able to amplify the innate immune response of human keratinocytes to pathogens by induction of AMP expression (Wanke, Steffen et al. 2011). Our SE_ΔΔΔ_ auxotroph promoted the expression hβD-2, a key AMP in the skin that has been shown to inhibit *S. aureus*, demonstrating its ability to favorably modulate host innate immunity (Sharma and Nagaraj 2015).

In this study, we describe the engineering of a live skin commensal, *S. epidermidis*, and characterize its favorable properties which support its potential use as an LBP for skin diseases. Properties conferred by the D-alanine auxtrophy, and the strain background itself, support the potential of SE_ΔΔΔ_ to serve either as an LBP itself, or as a recombinant host for expressing novel skin therapeutics. Future studies will evaluate the effect of this LBP candidate on human skin in the clinical setting.

## REFERENCES

Ali, S. M. and G. Yosipovitch (2013). “Skin pH: from basic science to basic skin care.” Acta Derm Venereol 93(3): 261–267.

Barbagelata, M. S., L. Alvarez, M. Gordiola, L. Tuchscherr, C. von Eiff, K. Becker, D. Sordelli and F. Buzzola (2011). “Auxotrophic mutant of Staphylococcus aureus interferes with nasal colonization by the wild type.” Microbes Infect 13(12-13): 1081–1090.

Bastos, M. C. F., H. Ceotto, M. L. V. Coelho and J. S. Nascimento (2009). “Staphylococcal Antimicrobial Peptides: Relevant Properties and Potential Biotechnological Applications.” Current Pharmaceutical Biotechnology 10(1): 38–61.

Bitschar, K., C. Wolz, B. Krismer, A. Peschel and B. Schittek (2017). “Keratinocytes as sensors and central players in the immune defense against Staphylococcus aureus in the skin.” J Dermatol Sci 87(3): 215–220.

Bose, J. L., P. D. Fey and K. W. Bayles (2013). “Genetic tools to enhance the study of gene function and regulation in Staphylococcus aureus.” Appl Environ Microbiol 79(7): 2218–2224.

Brogden, K. A. (2005). “Antimicrobial peptides: pore formers or metabolic inhibitors in bacteria?” Nat Rev Microbiol 3(3): 238–250.

Chen, C., V. Krishnan, K. Macon, K. Manne, S. V. Narayana and O. Schneewind (2013). “Secreted proteases control autolysin-mediated biofilm growth of Staphylococcus aureus.” J Biol Chem 288(41): 29440–29452.

Clausen, M. L. and T. Agner (2016). “Antimicrobial Peptides, Infections and the Skin Barrier.” Curr Probl Dermatol 49: 38–46.

Cogen, A. L., K. Yamasaki, J. Muto, K. M. Sanchez, L. Crotty Alexander, J. Tanios, Y. Lai, J. E. Kim, V. Nizet and R. L. Gallo (2010). “Staphylococcus epidermidis antimicrobial delta-toxin (phenol-soluble modulin-gamma) cooperates with host antimicrobial peptides to kill group A Streptococcus.” PLoS One 5(1): e8557.

Cogen, A. L., K. Yamasaki, K. M. Sanchez, R. A. Dorschner, Y. Lai, D. T. MacLeod, J. W. Torpey, M. Otto, V. Nizet, J. E. Kim and R. L. Gallo (2010). “Selective antimicrobial action is provided by phenol-soluble modulins derived from Staphylococcus epidermidis, a normal resident of the skin.” J Invest Dermatol 130(1): 192–200.

Conlan, S., L. A. Mijares, J. Becker, R. W. Blakesley, G. G. Bouffard, S. Brooks, H. Coleman, J. Gupta, N. Gurson, M. Park, B. Schmidt, P. J. Thomas, M. Otto, H. H. Kong, P. R. Murray and J. A. Segre (2012). “Staphylococcus epidermidis pan-genome sequence analysis reveals diversity of skin commensal and hospital infection-associated isolates.” Genome Biol 13(7): R64.

Conlan, S., L. A. Mijares, N. C. S. Program, J. Becker, R. W. Blakesley, G. G. Bouffard, S. Brooks, H. Coleman, J. Gupta, N. Gurson, M. Park, B. Schmidt, P. J. Thomas, M. Otto, H. H. Kong, P. R. Murray and J. A. Segre (2012). “Staphylococcus epidermidis pan-genome sequence analysis reveals diversity of skin commensal and hospital infection-associated isolates.” Genome Biology 13(7): R64–R64.

Diaz Heijtz, R., S. Wang, F. Anuar, Y. Qian, B. Bjorkholm, A. Samuelsson, M. L. Hibberd, H. Forssberg and S. Pettersson (2011). “Normal gut microbiota modulates brain development and behavior.” Proc Natl Acad Sci U S A 108(7): 3047–3052.

Dinulos, J. G. H., L. Mentele, L. P. Fredericks, B. A. Dale and G. L. Darmstadt (2003). “Keratinocyte expression of human beta defensin 2 following bacterial infection: role in cutaneous host defense.” Clinical and diagnostic laboratory immunology 10(1): 161–166.

Dore, J., M. C. Multon and J. M. Behier (2017). “The human gut microbiome as source of innovation for health: Which physiological and therapeutic outcomes could we expect?” Therapie 72(1): 21–38.

Dreher-Lesnick, S. M., S. Stibitz and P. E. Carlson, Jr. (2017). “U.S. Regulatory Considerations for Development of Live Biotherapeutic Products as Drugs.” Microbiol Spectr 5(5).

Dubin, G., D. Chmiel, P. Mak, M. Rakwalska, M. Rzychon and A. Dubin (2001). Molecular Cloning and Biochemical Characterisation of Proteases from Staphylococcus epidermidis. Biological Chemistry. 382:1575.

Dul, M. J. and F. E. Young (1973). “Genetic mapping of a mutant defective in D,L-alanine racemase in Bacillus subtilis 168.” J Bacteriol 115(3): 1212–1214.

Galac, M. R., J. Stam, R. Maybank, M. Hinkle, D. Mack, H. Rohde, A. L. Roth and P. D. Fey (2017). “Complete Genome Sequence of Staphylococcus epidermidis 1457.” Genome Announc 5(22).

Grice, E. A. (2014). “The skin microbiome: potential for novel diagnostic and therapeutic approaches to cutaneous disease.” Semin Cutan Med Surg 33(2): 98–103.

Grice, E. A. and J. A. Segre (2011). “The skin microbiome.” Nat Rev Microbiol 9(4): 244–253.

Iwase, T., Y. Uehara, H. Shinji, A. Tajima, H. Seo, K. Takada, T. Agata and Y. Mizunoe (2010). “Staphylococcus epidermidis Esp inhibits Staphylococcus aureus biofilm formation and nasal colonization.” Nature 465(7296): 346–349.

Kau, A. L., P. P. Ahern, N. W. Griffin, A. L. Goodman and J. I. Gordon (2011). “Human nutrition, the gut microbiome and the immune system.” Nature 474(7351): 327–336.

Kord, M., A. Ardebili, M. Jamalan, R. Jahanbakhsh, N. Behnampour and E. A. Ghaemi (2018). “Evaluation of Biofilm Formation and Presence of Ica Genes in Staphylococcus epidermidis Clinical Isolates.” Osong public health and research perspectives 9(4): 160–166.

Lai, Y., A. L. Cogen, K. A. Radek, H. J. Park, D. T. Macleod, A. Leichtle, A. F. Ryan, A. Di Nardo and R. L. Gallo (2010). “Activation of TLR2 by a small molecule produced by Staphylococcus epidermidis increases antimicrobial defense against bacterial skin infections.” J Invest Dermatol 130(9): 2211–2221.

Lai, Y., A. Di Nardo, T. Nakatsuji, A. Leichtle, Y. Yang, A. L. Cogen, Z. R. Wu, L. V. Hooper, R. R. Schmidt, S. von Aulock, K. A. Radek, C. M. Huang, A. F. Ryan and R. L. Gallo (2009). “Commensal bacteria regulate Toll-like receptor 3-dependent inflammation after skin injury.” Nat Med 15(12): 1377–1382.

Lannergård, J., C. von Eiff, G. Sander, T. Cordes, J. Seggewiβ, G. Peters, R. A. Proctor, K. Becker and D. Hughes (2008). “Identification of the Genetic Basis for Clinical Menadione-Auxotrophic Small-Colony Variant Isolates of &lt;em&gt;Staphylococcus aureus&lt;/em&gt.” Antimicrobial Agents and Chemotherapy 52(11): 4017.

Lee, J. and M. Blaber (2013). “Increased Functional Half-life of Fibroblast Growth Factor-1 by Recovering a Vestigial Disulfide Bond.” 2013 1(2).

Lee, J., S. Blaber, V. Dubey and M. Blaber (2011). A Polypeptide “Building Block” for the β-Trefoil Fold Identified by “Top-Down Symmetric Deconstruction”.

Li, C., F. Sun, H. Cho, V. Yelavarthi, C. Sohn, C. He, O. Schneewind and T. Bae (2010). “CcpA mediates proline auxotrophy and is required for Staphylococcus aureus pathogenesis.” J Bacteriol 192(15): 3883–3892.

Li, D., H. Lei, Z. Li, H. Li, Y. Wang and Y. Lai (2013). “A novel lipopeptide from skin commensal activates TLR2/CD36-p38 MAPK signaling to increase antibacterial defense against bacterial infection.” PLoS One 8(3): e58288.

Linehan, J. L., O. J. Harrison, S. J. Han, A. L. Byrd, I. Vujkovic-Cvijin, A. V. Villarino, S. K. Sen, J. Shaik, M. Smelkinson, S. Tamoutounour, N. Collins, N. Bouladoux, A. Dzutsev, S. P. Rosshart, J. H. Arbuckle, C. R. Wang, T. M. Kristie, B. Rehermann, G. Trinchieri, J. M. Brenchley, J. J. O’Shea and Y. Belkaid (2018). “Non-classical Immunity Controls Microbiota Impact on Skin Immunity and Tissue Repair.” Cell 172(4): 784–796.e718.

Livak, K. J. and T. D. Schmittgen (2001). “Analysis of relative gene expression data using real-time quantitative PCR and the 2(-Delta Delta C(T)) Method.” Methods 25(4): 402–408.

Mack, D., N. Siemssen and R. Laufs (1992). “Parallel induction by glucose of adherence and a polysaccharide antigen specific for plastic-adherent Staphylococcus epidermidis: evidence for functional relation to intercellular adhesion.” Infection and immunity 60(5): 2048–2057.

MacLea, K. S. and A. M. Trachtenberg (2017). “Complete Genome Sequence of Staphylococcus epidermidis ATCC 12228 Chromosome and Plasmids, Generated by Long-Read Sequencing.” Genome Announc 5(36).

Moon, J., L., A. Banbula, A. Oleksy, J. Mayo, A. and J. Travis (2001). Isolation and Characterization of a Highly Specific Serine Endopeptidase from an Oral Strain of Staphylococcus epidermidis. Biological Chemistry. 382:1095.

Moscoso, M., P. Garcia, M. P. Cabral, C. Rumbo and G. Bou (2018). “A D-Alanine auxotrophic live vaccine is effective against lethal infection caused by Staphylococcus aureus.” Virulence 9(1): 604–620.

Mulcahy, M. E., J. A. Geoghegan, I. R. Monk, K. M. O’Keeffe, E. J. Walsh, T. J. Foster and R. M. McLoughlin (2012). “Nasal colonisation by Staphylococcus aureus depends upon clumping factor B binding to the squamous epithelial cell envelope protein loricrin.” PLoS Pathog 8(12): e1003092.

Naik, S., N. Bouladoux, J. L. Linehan, S. J. Han, O. J. Harrison, C. Wilhelm, S. Conlan, S. Himmelfarb, A. L. Byrd, C. Deming, M. Quinones, J. M. Brenchley, H. H. Kong, R. Tussiwand, K. M. Murphy, M. Merad, J. A. Segre and Y. Belkaid (2015). “Commensal-dendritic-cell interaction specifies a unique protective skin immune signature.” Nature 520(7545): 104–108.

Naik, S., N. Bouladoux, C. Wilhelm, M. J. Molloy, R. Salcedo, W. Kastenmuller, C. Deming, M. Quinones, L. Koo, S. Conlan, S. Spencer, J. A. Hall, A. Dzutsev, H. Kong, D. J. Campbell, G. Trinchieri, J. A. Segre and Y. Belkaid (2012). “Compartmentalized control of skin immunity by resident commensals.” Science 337(6098): 1115–1119.

Nakatsuji, T., T. H. Chen, A. M. Butcher, L. L. Trzoss, S.-J. Nam, K. T. Shirakawa, W. Zhou, J. Oh, M. Otto, W. Fenical and R. L. Gallo (2018). “A commensal strain of &lt;em&gt;Staphylococcus epidermidis&lt;/em&gt; protects against skin neoplasia.” Science Advances 4(2): eaao4502.

Nakatsuji, T., T. H. Chen, A. M. Butcher, L. L. Trzoss, S. J. Nam, K. T. Shirakawa, W. Zhou, J. Oh, M. Otto, W. Fenical and R. L. Gallo (2018). “A commensal strain of Staphylococcus epidermidis protects against skin neoplasia.” Sci Adv 4(2): eaao4502.

Nakatsuji, T., T. H. Chen, S. Narala, K. A. Chun, A. M. Two, T. Yun, F. Shafiq, P. F. Kotol, A. Bouslimani, A. V. Melnik, H. Latif, J.-N. Kim, A. Lockhart, K. Artis, G. David, P. Taylor, J. Streib, P. C. Dorrestein, A. Grier, S. R. Gill, K. Zengler, T. R. Hata, D. Y. M. Leung and R. L. Gallo (2017). “Antimicrobials from human skin commensal bacteria protect against *Staphylococcus aureus* and are deficient in atopic dermatitis.” Science Translational Medicine 9(378).

Neopane, P., H. P. Nepal, R. Shrestha, O. Uehara and Y. Abiko (2018). “In vitro biofilm formation by Staphylococcus aureus isolated from wounds of hospital-admitted patients and their association with antimicrobial resistance.” Int J Gen Med 11: 25–32.

Niyonsaba, F., C. Kiatsurayanon, P. Chieosilapatham and H. Ogawa (2017). “Friends or Foes? Host defense (antimicrobial) peptides and proteins in human skin diseases.” Exp Dermatol 26(11): 989–998.

Nodake, Y., S. Matsumoto, R. Miura, H. Honda, G. Ishibashi, S. Matsumoto, I. Dekio and R. Sakakibara (2015). “Pilot study on novel skin care method by augmentation with Staphylococcus epidermidis, an autologous skin microbe--A blinded randomized clinical trial.” J Dermatol Sci 79(2): 119–126.

Nodake, Y., S. Matsumoto, R. Miura, H. Honda, G. Ishibashi, S. Matsumoto, I. Dekio and R. Sakakibara (2015). “Pilot study on novel skin care method by augmentation with Staphylococcus epidermidis, an autologous skin microbe--A blinded randomized clinical trial.” Journal of dermatological science 79(2): 119–126.

Oh, J., A. L. Byrd, C. Deming, S. Conlan, H. H. Kong and J. A. Segre (2014). “Biogeography and individuality shape function in the human skin metagenome.” Nature 514(7520): 59–64.

Oh, J., A. F. Freeman, M. Park, R. Sokolic, F. Candotti, S. M. Holland, J. A. Segre and H. H. Kong (2013). “The altered landscape of the human skin microbiome in patients with primary immunodeficiencies.” Genome Res 23(12): 2103–2114.

Pasparakis, M., I. Haase and F. O. Nestle (2014). “Mechanisms regulating skin immunity and inflammation.” Nat Rev Immunol 14(5): 289–301.

Pfaffl, M. W. (2001). “A new mathematical model for relative quantification in real-time RT-PCR.” Nucleic Acids Res 29(9): e45.

Pivarcsi, A., L. Kemény and A. Dobozy (2004). “Innate Immune Functions of the Keratinocytes.” Acta Microbiologica et Immunologica Hungarica 51(3): 303–310.

Powers, C. E., D. B. McShane, P. H. Gilligan, C. N. Burkhart and D. S. Morrell (2015). “Microbiome and pediatric atopic dermatitis.” J Dermatol.

Ravel, J., P. Gajer, Z. Abdo, G. M. Schneider, S. S. Koenig, S. L. McCulle, S. Karlebach, R. Gorle, J. Russell, C. O. Tacket, R. M. Brotman, C. C. Davis, K. Ault, L. Peralta and L. J. Forney (2011). “Vaginal microbiome of reproductive-age women.” Proc Natl Acad Sci U S A 108 Suppl 1: 4680–4687.

Sahl, H. G., Bierbaum, G. (1998). “LANTIBIOTICS: Biosynthesis and Biological Activities of Uniquely Modified Peptides from Gram-Positive Bacteria.” Annual Review of Microbiology 52(1): 41–79.

Salzman, N. H., K. Hung, D. Haribhai, H. Chu, J. Karlsson-Sjoberg, E. Amir, P. Teggatz, M. Barman, M. Hayward, D. Eastwood, M. Stoel, Y. Zhou, E. Sodergren, G. M. Weinstock, C. L. Bevins, C. B. Williams and N. A. Bos (2010). “Enteric defensins are essential regulators of intestinal microbial ecology.” Nat Immunol 11(1): 76–82.

Schittek, B., M. Paulmann, I. Senyurek and H. Steffen (2008). “The role of antimicrobial peptides in human skin and in skin infectious diseases.” Infect Disord Drug Targets 8(3): 135–143.

Sharma, H. and R. Nagaraj (2015). “Human β-defensin 4 with non-native disulfide bridges exhibit antimicrobial activity.” PloS one 10(3): e0119525–e0119525.

Simanski, M., R. Glaser and J. Harder (2014). “Human skin engages different epidermal layers to provide distinct innate defense mechanisms.” Exp Dermatol 23(4): 230–231.

Stacy, A. and Y. Belkaid (2019). “Microbial guardians of skin health.” Science 363(6424): 227–228.

Sugimoto, S., T. Iwamoto, K. Takada, K. Okuda, A. Tajima, T. Iwase and Y. Mizunoe (2013). “Staphylococcus epidermidis Esp degrades specific proteins associated with Staphylococcus aureus biofilm formation and host-pathogen interaction.” J Bacteriol 195(8): 1645–1655.

Thompson, R. J., H. G. Bouwer, D. A. Portnoy and F. R. Frankel (1998). “Pathogenicity and Immunogenicity of a Listeria monocytogenes Strain That Requires d-Alanine for Growth.” Infection and Immunity 66(8): 3552–3561.

Vandecandelaere, I., P. Depuydt, H. J. Nelis and T. Coenye (2014). “Protease production by Staphylococcus epidermidis and its effect on Staphylococcus aureus biofilms.” Pathog Dis 70(3): 321–331.

Vemuri, R. C., R. Gundamaraju, T. Shinde and R. Eri (2017). “Therapeutic interventions for gut dysbiosis and related disorders in the elderly: antibiotics, probiotics or faecal microbiota transplantation?” Benef Microbes 8(2): 179–192.

Wan, P. and J. Chen (2020). “A Calm, Dispassionate Look at Skin Microbiota in Atopic Dermatitis: An Integrative Literature Review.” Dermatol Ther (Heidelb) 10(1): 53–61.

Wang, D., P. W. L. Tai and G. Gao (2019). “Adeno-associated virus vector as a platform for gene therapy delivery.” Nature Reviews Drug Discovery 18(5): 358–378.

Wanke, I., H. Steffen, C. Christ, B. Krismer, F. Gotz, A. Peschel, M. Schaller and B. Schittek (2011). “Skin commensals amplify the innate immune response to pathogens by activation of distinct signaling pathways.” J Invest Dermatol 131(2): 382–390.

Wei, W., Z. Cao, Y. L. Zhu, X. Wang, G. Ding, H. Xu, P. Jia, D. Qu, A. Danchin and Y. Li (2006). “Conserved genes in a path from commensalism to pathogenicity: comparative phylogenetic profiles of Staphylococcus epidermidis RP62A and ATCC12228.” BMC Genomics 7: 112.

Wei, W., J. Wiggins, D. Hu, V. Vrbanac, D. Bowder, M. Mellon, A. Tager, J. Sodroski and S. H. Xiang (2019). “Blocking HIV-1 Infection by Chromosomal Integrative Expression of Human CD4 on the Surface of Lactobacillus acidophilus ATCC 4356.” J Virol 93(8).

Weinstein, M. P. and J. S. Lewis (2020). “The Clinical and Laboratory Standards Institute Subcommittee on Antimicrobial Susceptibility Testing: Background, Organization, Functions, and Processes.” Journal of Clinical Microbiology 58(3): e01864–01819.

Weyrich, L. S., S. Dixit, A. G. Farrer, A. J. Cooper and A. J. Cooper (2015). “The skin microbiome: Associations between altered microbial communities and disease.” Australas J Dermatol.

Wikoff, W. R., A. T. Anfora, J. Liu, P. G. Schultz, S. A. Lesley, E. C. Peters and G. Siuzdak (2009). “Metabolomics analysis reveals large effects of gut microflora on mammalian blood metabolites.” Proc Natl Acad Sci U S A 106(10): 3698–3703.

Yamasaki, K. and R. L. Gallo (2008). “Antimicrobial peptides in human skin disease.” European journal of dermatology: EJD 18(1): 11–21.

Yin, X., J. Goudriaan, E. A. Lantinga, J. Vos and H. J. Spiertz (2003). “A flexible sigmoid function of determinate growth.” Ann Bot 91(3): 361–371.

Zhang, Y. Q., S. X. Ren, H. L. Li, Y. X. Wang, G. Fu, J. Yang, Z. Q. Qin, Y. G. Miao, W. Y. Wang, R. S. Chen, Y. Shen, Z. Chen, Z. H. Yuan, G. P. Zhao, D. Qu, A. Danchin and Y. M. Wen (2003). “Genome-based analysis of virulence genes in a non-biofilm-forming Staphylococcus epidermidis strain (ATCC 12228).” Mol Microbiol 49(6): 1577–1593.

