## Supplemental Information for "Controlling the Growth of the Skin Commensal *Staphylococcus epidermidis* Using D-Alanine Auxotrophy"

**This PDF file includes:**

Supplementary Table S1

Supplementary Table S2

Supplementary Table S3

Supplementary Figure S1

**SUPPLEMENTARY TABLES**

**Supplementary Table S1. Strains and plasmids used or developed in this study**

| **Strain or plasmid** | **Relevant characteristics** | **Source or reference** |
| --- | --- | --- |
| *Escherichia coli* | | |
| Top10 |  | Life Technologies, Inc., Carlsbad, CA, USA |
| GM2163 (*dam^-^/dcm^-^)* | Adenine and cytosine methylation-deficient *E. coli* | Source?New England Biolabs, Ipswich, MA, USA |
| *Staphylococcus epidermidis* |  |  |
| NRRL B-4268 | Non-biofilm-forming S. epidermidis strain. Originally deposited as PCI 1200. Re-deposited as ATCC 12228, then re-deposited as NRRL B-4268. | USDA ARS NRRL Collection  (Zhang, Ren et al. 2003) |
| SEΔ*alr1*Δ*alr2* | Alanine racemase-deficient NRRL B-4268 | This study |
| SEΔ*alr1*Δ*alr2*Δ*dat*, or SE_ΔΔΔ_ | d-alanine auxotrophic NRRL B-4268 | This study |
| 1457 | *Ica -* positive, biofilm-forming | (Galac, Stam et al. 2017) |
| *Staphylococcus aureus* |  |  |
| RN4220 | Restriction-deficient 8325-4 | (Kreiswirth, Lofdahl et al. 1983) |
| Plasmids |  |  |
| pJB38 | Allelic exchange shuttle vector, Cam^r^/ Amp^r^ | (Cheung, Bayer et al. 2004) |
| pJB38-1674KO | pJB38 containing the 5’ upstream and 3’ downstream flanking sequences of the target gene SE1674 | This study |
| pJB38-1079KO | pJB38 containing the 5’ upstream and 3’ downstream flanking sequences of SE1079 | This study |
| pJB38-1423KO | pJB38 containing the 5’ upstream and 3’ downstream flanking sequences of SE1423 | This study |

**Supplementary Table S2. Primers Used in This Study**

| **Name** | **Sequence (5’ to 3’)** | **Application** |
| --- | --- | --- |
| 1674-5F (SalI) | atgc**gtcgac**TTGGTACATGAAAGGTGATAC | To amplify the 5’ flanking region of SE1674 (1.0 Kb) |
| 1674-5R | caaatttcctaatcagtgactataATATATGTCCTCCTTGAAACTACTTAC |  |
| 1674-3F | gtaagtagtttcaaggaggacatatatTATAGTCACTGATTAGGAAATTTG | To amplify the 3’ flanking region of SE1674 (1.0 Kb) |
| 1674-3R (EcoRI) | acgt**gaattc**TTCCACGAAATGCGCCTC |  |
| 1674-F | ATGTCAGAGAAGTTTTATAGAG | To amplify the entire SE1674 CDS (1.2 Kb, deleted in 1674 KO strains) |
| 1674-R | CTATTTTAACAATTCGTTAGTAAC |  |
| JB-Cm-F | TTGATTTAGACAATTGGAAGAG | To amplify part of the chloramphenicol selection marker (0.7 Kb) in pJB38 |
| JB-Cm-R | AAGTACAGTCGGCATTATCTC |  |
| 1079-5F (EcoRI) | acgt**gaattc**GTTACATTGCACAGAAG | To amplify 5’ flanking region of SE1079 (1.2 Kb) |
| 1079-5R | ctccttcataagagaatcgtgTTGCTTTACACCTCTTTATAATTTC |  |
| 1079-3F | gaaattataaagaggtgtaaagcaaCACGATTCTCTTATGAAGGAG | To amplify 3’ flanking region of SE1079 (1.0 Kb) |
| 1079-3R (SalI) | acgt**gtcgac**ACGCCTCATACTGTGCACCATAAAG |  |
| 1079-F | TTGACAGCAATTTGGTCATTAG | To amplify SE1079 CDS (1.1 Kb, deleted in 1079 KO strains) |
| 1079-R | CTCCTTCATAAGAGAATCGTG |  |
| 1423-5F  (EcoRI) | atgc**gaattc**ATGAGCGATACTTATTTGAATC | Amplification of 5’ flanking region of SE1423 (0.5 Kb) |
| 1423-5R | ctatgcgattgaatatacttttcCTTAGCATCCTCTTCATTAAC |  |
| 1423-3F | gttaatgaagaggatgctaaggaAAAGTATATTCAATCGCATAG | Amplification of 3’ flanking region of SE1423 (1.0 Kb) |
| 1423-3R  (SalI) | agct**gtcgac**AGCAGCATACCAATGTCAATC |  |
| 1423-F | CATACGAAGATCGAGGCTAC | Amplification of a partial SE1423 (0.7 Kb) |
| 1423-R | GTACCAACTTGTCCGTCTTG |  |
| DEFB4A | Hs00175474_m1 (TAQMANGene Expression Assays, Applied Biosystems) | Amplification of Beta 4 Defensin |
| S100A7 | Hs00161488_m1 (TAQMANGene Expression Assays, Applied Biosystems) | Amplification of S100 Calcium-binding protein A7 |
| B2M | Hs_00984230_m1 (TAQMANGene Expression Assays, Applied Biosystems) | Amplification of B2 microglobulin, a housekeeping gene |

**Supplementary Table S3. Growth and survival of SE_ΔΔΔ_ in defibrinated pooled healthy human blood after 24hr incubation**

| **Inoculum at T=0 (CFU/mL)** | **CFU/mL after 24 hr growth** | |
| --- | --- | --- |
|  | SE_ΔΔΔ +_ d-alanine | SE_ΔΔΔ -_ d-alanine |
| 1.0 x 10^0^ | 0 x 10^0^ | 0 |
| 6.0 x 10^2^ | 1.0 x 10^5^ | 0 |
| 2.0 x 10^4^ | 2.0 x 107 | 0 |
| 4.0 x 10^6^ | 1.0 x 10^8^ | 0 |
| 1.4 x 10^8^ | 2.0 x 10^8^ | 1.3 x 10^8^ |

The growth of SE_ΔΔΔ_ was tested in blood +/- 100 µg/mL of d-alanine. Results are the averages of two experimental replicates

**Supplementary Figure S1. Frequency of spontaneous reversion of D-alanine auxotrophy in SE_ΔΔΔ_**

Condition: No D-alanine	100 µg/mL D-alanine	8 ng/mL rifampicin/

100 µg/mL D-alanine

Frequency: **<4.8 x 10^-11^	Confluent Growth	6 x 10^-8^**

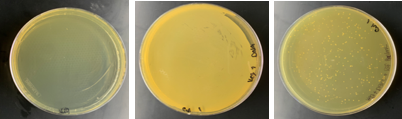

Vegitone agar plates were inoculated with 2.1 x 10^10^ CFU of SE_ΔΔΔ._ and grown at 37 °C for 24 hours (A), without D-alanine, (B), with 100 µg/mL D-alanine, and (C), Vegitone agar supplemented with 100 µg/mL D-alanine and 8 ng/mL of rifampicin (positive control). A representative result of 4 independent trials is shown. The frequency of spontaneous resistance against rifampicin observed here is consistent with literature reports for this antibiotic.

**REFERENCES**

Cheung, A. L., A. S. Bayer, G. Zhang, H. Gresham and Y. Q. Xiong (2004). "Regulation of virulence determinants in vitro and in vivo in Staphylococcus aureus." FEMS Immunol Med Microbiol **40**(1): 1-9.

Galac, M. R., J. Stam, R. Maybank, M. Hinkle, D. Mack, H. Rohde, A. L. Roth and P. D. Fey (2017). "Complete Genome Sequence of Staphylococcus epidermidis 1457." Genome Announc **5**(22).

Kreiswirth, B. N., S. Lofdahl, M. J. Betley, M. O'Reilly, P. M. Schlievert, M. S. Bergdoll and R. P. Novick (1983). "The toxic shock syndrome exotoxin structural gene is not detectably transmitted by a prophage." Nature **305**(5936): 709-712.

Zhang, Y. Q., S. X. Ren, H. L. Li, Y. X. Wang, G. Fu, J. Yang, Z. Q. Qin, Y. G. Miao, W. Y. Wang, R. S. Chen, Y. Shen, Z. Chen, Z. H. Yuan, G. P. Zhao, D. Qu, A. Danchin and Y. M. Wen (2003). "Genome-based analysis of virulence genes in a non-biofilm-forming Staphylococcus epidermidis strain (ATCC 12228)." Mol Microbiol **49**(6): 1577-1593.
